# The stress of carrying CRISPR-Cas

**DOI:** 10.1101/2024.04.03.587888

**Authors:** Dorina Haider, Richard Bauer, Aline Grempels, Rebecca Roscher, Cagil Ceren Aslan, Stefanie Mauerer, Barbara Spellerberg

## Abstract

*Streptococcus anginosus* (*S. anginosus*) is a commensal that can cause severe invasive bacterial infections. A considerable percentage of *S. anginosus* strains harbor CRISPR-Cas systems, which apart from being a bacterial immunity system can play an important role regarding the adaptation to environmental stress. The functionality of *S. anginosus* CRISPR-Cas systems has previously not been investigated. To address this, we created a set of deletion mutants in the CRISPR-Cas type II-A system of the *S. anginosus* SK52 type strain, targeting the nuclease Cas9 and the CRISPR array. Testing these strains in a plasmid clearance assay, we were able to confirm CRISPR-Cas activity. Furthermore, the role of the *S. anginosus* CRISPR-Cas system was investigated under various stress conditions such as UV light, hydrogen peroxide exposure, and high-temperatures in wildtype *S. anginosus* and CRISPR-Cas mutant strains. Under these conditions, survival was significantly lower in strains carrying *cas9.* Bacterial growth and metabolic activity in Alamar blue assays was also negatively affected by the presence of *cas9* in *S. anginosus*. In summary we found that the presence of a functional CRISPR-Cas system in *S. anginosus* leads to measurable metabolic and fitness costs for the wildtype strain. Carrying *cas9* was associated with an impaired stress response in our experiments and may thus explain, why many strains of this species lack CRISPR-Cas.

**Author Summary:** The bacterial immunity system CRIPRS-Cas provides protection against invading foreign genetic material. Despite this obvious advantage only about 50% of bacteria carry CRISPR-Cas. To investigate the CRISPR system of *Streptococcus anginosus*, which can cause serious bacterial infections and has recently been linked to gastric cancer, we created a set of mutants in different loci of the CRISPR system. Exposing these mutants to stress through UV-light, hydrogen peroxide and high temperatures, we could show that carrying the CRISPR nuclease gene Cas9 is associated with impaired survival under harsh conditions. Strains lacking the nuclease gene had a better growth and higher metabolic activity than the wildtype strain. In summary we found that the presence of a functional CRISPR-Cas system in *S. anginosus* leads to considerable metabolic and fitness costs. Carrying *cas9* was associated with an impaired stress response in our experiments and may thus explain, why many strains of this species lack CRISPR-Cas.

## Introduction

Prokaryotic organisms have evolved strategies to prevent infections by bacteriophages or other invading genetic elements (1). These include the CRISPR systems (Clustered Regularly Interspaced Short Palindromic Repeats) containing distinct CRISPR-associated (*cas*) genes and a CRISPR array composed of unique spacer sequences interspersed with short repeats (1–5).

To adapt to novel infections, the range of defense is continuously expanded by the integration of new spacers derived from foreign genetic elements into the CRISPR array. These are transcribed and processed into CRISPR RNAs (crRNAs), consisting of repeat- and spacer sequences (6–9). Mature crRNAs form complexes with Cas proteins and mediate sequence-specific cleavage of target motifs on invading nucleic acids (10–12). Due to their great diversity, CRISPR-Cas systems are divided into two classes including three main types each and several subtypes based upon *cas* gene content, repeat sequence and the organization of the CRISPR loci (class 1 comprising type I, III and IV; class 2 comprising type II, V, VI) (4,7,13). While class 1 systems employ multi-subunit Cas protein complexes, the effector modules of class 2 systems only contain a single Cas protein as the associated type II system with its prominent signature gene *cas9* (4,14,15). The simplicity of the effector module Cas9 has led to the extensive usage of *cas9* genes for genetic manipulation in a multitude of different organisms (16,17). Despite the obvious advantages of carrying CRISPR-Cas it can only be detected in about 50 % of bacterial isolates (18). The reason for the absence in a large proportion of strains or even entire species like *Streptococcus pneumoniae* remain elusive (19).

Besides the canonical immunity functions of CRISPR-Cas, non-canonical functions have gained more attention in recent years. There is increasing evidence that CRISPR-Cas may regulate bacterial physiology, more precisely is involved in DNA repair, regulation of endogenous gene expression and bacterial virulence (20–22). Furthermore, it is associated with stress tolerance towards a variety of environmental factors. As response to stress CRISPR-Cas systems are transcriptionally activated and have been shown to prevent stress-promoted damages (23,24). While all bacteria are exposed to a wide range of environmental stresses, human commensals and pathogens such as *Streptococcus anginosus* encounter a multitude of different microenvironments and stressors in the host (21).

*S. anginosus* is found primarily as a commensal of mucosal membranes colonizing many areas of the human body including the oral cavity, the gastrointestinal and the urogenital tract (25–28). It belongs to the *Streptococcus anginosus* group (SAG) together with the closely related species *Streptococcus constellatu*s and *Streptococcus intermedius* (29,30). During the last years, *S. anginosus* has been increasingly identified in invasive infections emphasizing its clinical importance as an emerging bacterial pathogen (31–33). It is frequently isolated from abscesses and blood cultures, the respiratory tract of cystic fibrosis patients and has recently been associated with gastric cancer (34–38). The incidence of SAG infections (8.65/100,000) even exceeds the combined incidence rates of group A and B streptococci, in population-based surveillance studies (31,39).

We previously reported on the diversity of *Streptococcus anginosus* CRISPR-Cas type II-A systems and the negative association of hemolysin genes with the carriage of CRISPR-Cas in *S. anginosus* (16,40). To get a deeper insight into the role of CRISPR-Cas beyond defense, we studied the stress response related to the presence of a functional CRISPR-Cas system in this species.

## Results

### Expression and functionality of CRISPR-Cas system in *S. anginosus*

While first genomes of SAG revealed that many *S. anginosus* strains carry a CRISPR-Cas system (43), its functionality has previously not been investigated. To address this, we created a set of mutants in the CRISPR-Cas type II-A system of *S. anginosus* SK52 (Figure S1). These comprise a *cas9* deletion (Δ*cas9*), CRISPR array deletion (ΔCRISPR) and *cas9* complementation strain (Δ*cas9*::*cas9*). The Δ*cas9* mutant carries a 590 bp deletion resulting in a frameshift and a 164 amino acid long truncated Cas9 protein. The ΔCRISPR mutant lacks the complete 1.600 bp CRISPR array and carries a single point mutation downstream of the CRISPR array acquired during mutant generation. In the Δ*cas9::cas9* strain *cas9* was complemented by chromosomal integration resulting in a 1.397 amino acid long native Cas9 protein (Figure S1). To evaluate CRISPR-Cas activity in these strains, we performed a plasmid clearance assay. The plasmid constructs were designed to contain a single native spacer present in the CRISPR locus of *S. anginosus* SK52, along with either a typical streptococcal PAM sequence (TGG) or a non-functional PAM (AAA). The vectors pAT28_SP1_TGG and pAT28_SP1_AAA were then transformed into *S. anginosus* wildtype and mutant strains. If the type II-A CRISPR-Cas system of *S. anginosus* SK52 is functional, successful plasmid transformation should be prevented or at least reduced in presence of a correct PAM through cleavage of the plasmid by Cas9. By using these plasmids, CRISPR interference activity could indeed be demonstrated (Figure 1). While pAT28_SP1_AAA and the empty vector pAT28 were transformable in all strains, pAT28_SP1_TGG was only transformable in the Δ*cas9* and the ΔCRISPR strain but not in the wildtype *S. anginosus* and the Δ*cas9::cas9* mutant.

**Figure 1:**
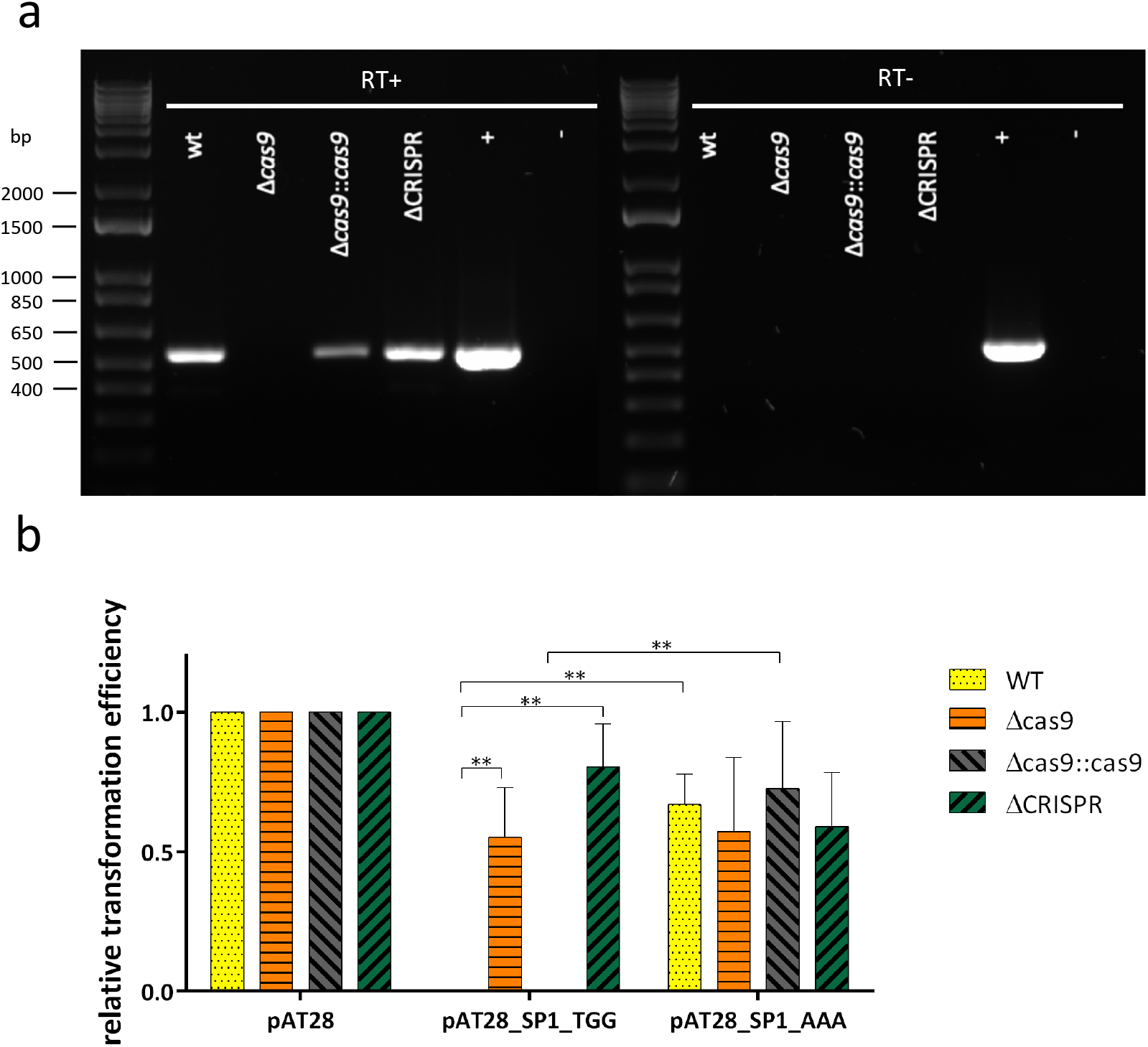
Expression and functionality of CRISPR-Cas system in *S. anginosus*. (a) RT-PCR performed with 20 ng isolated RNA of *S. anginosus* SK52 (WT), *cas9* deletion (Δ*cas9*), *cas9* complementation (Δ*cas9::cas9*) and CRISPR array deletion (ΔCRISPR) strain verifying expression of *cas9* using Primer 13/41. Complete DNase digestion was controlled by conducting PCR without RT step. Controls with genomic DNA of *S. anginosus* SK52 are indicated by “+” and negative controls without nucleic acid by “-“. The 1 kb Plus DNA Ladder (Invitrogen) served as molecular weight marker. (b) Confirming the functionality of *S. anginosus* CRISPR-Cas system by interference with plasmid transformation. Relative transformation efficiencies of created plasmids normalized to the empty vector pAT28 were obtained via competence-based transformation assay. Vectors were designed to contain an endogenous spacer of *S. anginosus* SK52 CRISPR-Cas system followed by a functional or non-functional PAM sequence. Mean values and standard deviations of six independent experiments are shown. Mann-Whitney-U test was performed to illustrate significant differences as indicated (**p < 0.01).

To further characterize CRISPR-Cas activity in *S. anginosus* SK52, an EGFP reporter plasmid harboring the *cas9* promoter was created as detailed in Materials and Methods. *Cas9* expression was determined by measuring EGFP activity in different growth phases and under various potential CRISPR-Cas inducing conditions (Figure S2 & Figure S3). In addition to the different growth stages, it was determined whether *cas9* expression is induced during stress mediated by antimicrobial substances as ampicillin and ciprofloxacin as well as hydrogen peroxide. Further, glucose supplementation in different concentrations as well as stimulation via DNA were tested. For these experiments CRISPR-Cas functionality assessing plasmids containing a single endogenous spacer of *S. anginosus* SK52 along with either a functional or non-functional PAM as well as chromosomal DNA of *S. anginosus* were supplemented in the presence or absence of CSP. While EGFP fluorescence was easily detectable, significant differences in *cas9* expression in *S. anginosus* SK52 could not be observed over different growth stages and were not induced as response to oxidative stress, glucose, or exposure to chromosomal DNA or CRISPR-Cas functionality assessing plasmids. Overall *cas9* expression appeared to take place at constant level over a range of different environmental growth conditions.

To confirm the expression of *cas9*, RT-PCR was performed using 20 ng isolated RNA of *S. anginosus*. Complete DNase digestion was controlled by additionally conducting the PCR without reverse transcription step. A clear and well reproducible expression of *cas9* RNA could be observed in the RT-experiments (Figure 1A) confirming the results of our reporter gene analysis and the plasmid interference assay. Comparing the *cas9* transcript level, a lower amount of transcript was detected in the Δ*cas9::cas9* strain in contrast to the wildtype and the ΔCRISPR mutant while expression was completely abolished in the Δ*cas9* strain. This data demonstrates to our knowledge for the first time a functional CRISPR-Cas system in the species *S. anginosus*. *Cas9* transcription was detectable in RT-PCR and translation was shown in EGFP reporter assays. However, conditions leading to a clear induction or repression of *cas9* transcription could not be defined, suggesting the presence of a more or less constant level of Cas9 expression in *S. anginosus*.

Role of *S. anginosus* SK52 CRISPR-Cas system under stress conditions

Various studies have demonstrated a role for CRISPR-Cas besides the immunity function. In *Streptococcus mutans* it has been shown that CRISPR-Cas systems can be activated as response to environmental changes in order to resist and combat stress and to promote bacterial survival (24,44). To study the stress tolerance of *S. anginosus* SK52, resistance against UV light and oxidative stressors as well as high-temperature was examined in wildtype and CRISPR-Cas mutant strains. We tested the influence of *cas9* expression under DNA-damaging conditions by investigating the survival of these strains under UV irradiation. Viability of the *cas9* deletion mutant was drastically enhanced when exposed to UV irradiation compared to the wildtype *S. anginosus* SK52, the Δ*cas9::cas9* and the ΔCRISPR mutant, suggesting that expression of *cas9* may affect the tolerance of UV mediated stress (Figure 2). The lethal irradiation time (defined as < 5 CFU) varied greatly among the wildtype and mutant strains. While Δ*cas9* nearly survived 90 sec of UV light exposure, the wildtype strain and the ΔCRISPR mutant almost reached 55 sec and the Δ*cas9::cas9* mutant around 50 sec. A similar picture was observed for survival in the presence of hydrogen peroxide or heat as stress conditions. Δ*cas9* was increased in its ability to tolerate stress induced by H_2_O_2_, suggesting the absence of *cas9* played a role in responding to extracellular oxidative stress (Figure 3B). Exposed to temperature stress at 50 °C, the *cas9* deficient mutant significantly survived this harsh condition best compared to the wildtype and the other mutant strains, suggesting that *S. anginosus* SK52 *cas9* has a role in temperature stress tolerance (Figure 3A). Taken together the presence of an intact *cas9* gene appeared to impair survival under different stress conditions in *S. anginosus*.

**Figure 2:**
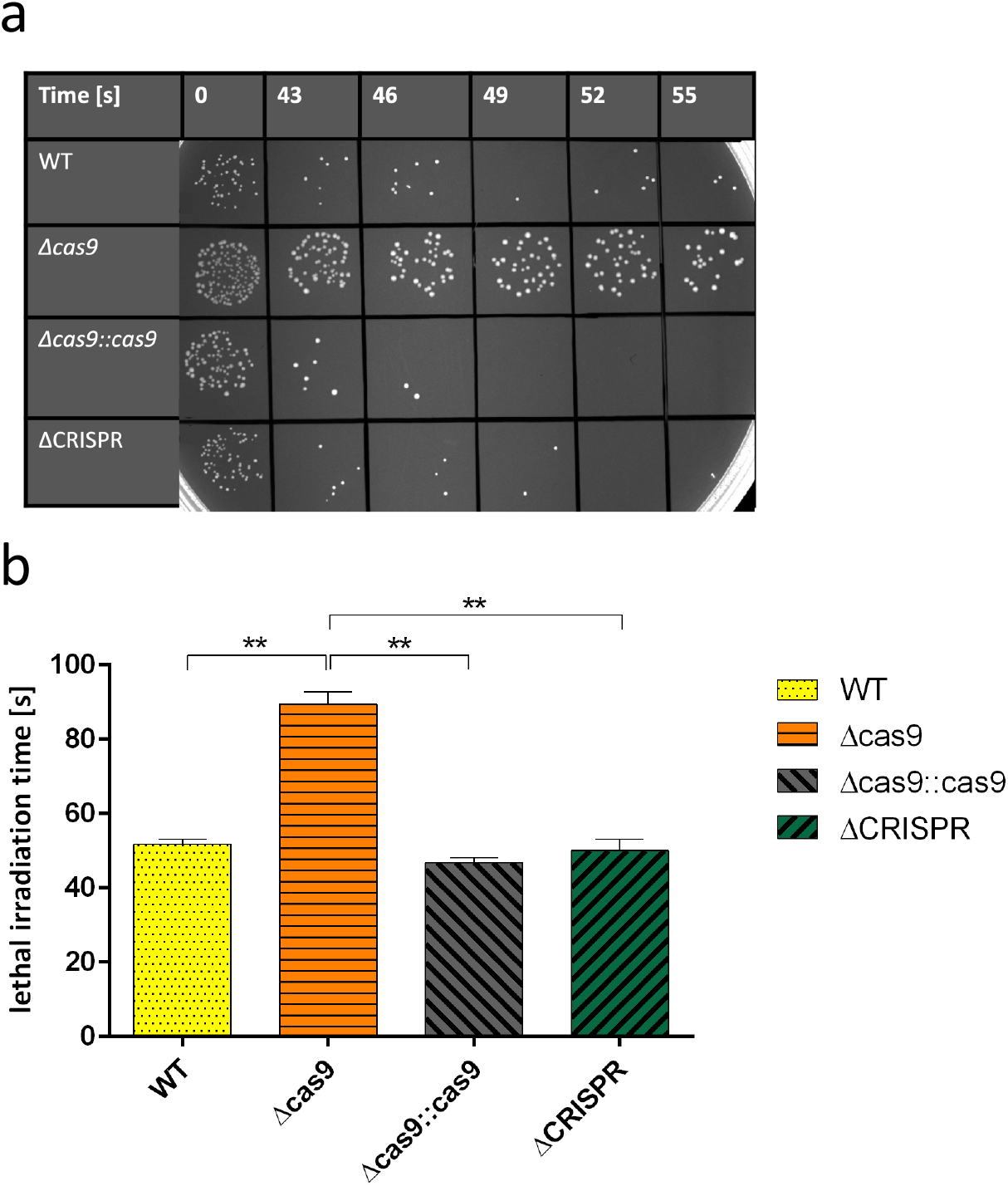
Effects of UV irradiation on viability of *S. anginosus* SK52 and mutant strains Δ*cas9*, Δ*cas9::cas9* and ΔCRISPR. (a) Strains on this agar plate were exposed to UV irradiation for 43 s – 96 s on THY agar followed by 24 h of growth at 37 °C. (b) Results are given as lethal irradiation time, where less than five CFU could be counted. Shown is the mean ± standard deviation of five independent experiments. Significant differences were calculated with Mann-Whitney-U test (**p < 0.01).

**Figure 3:**
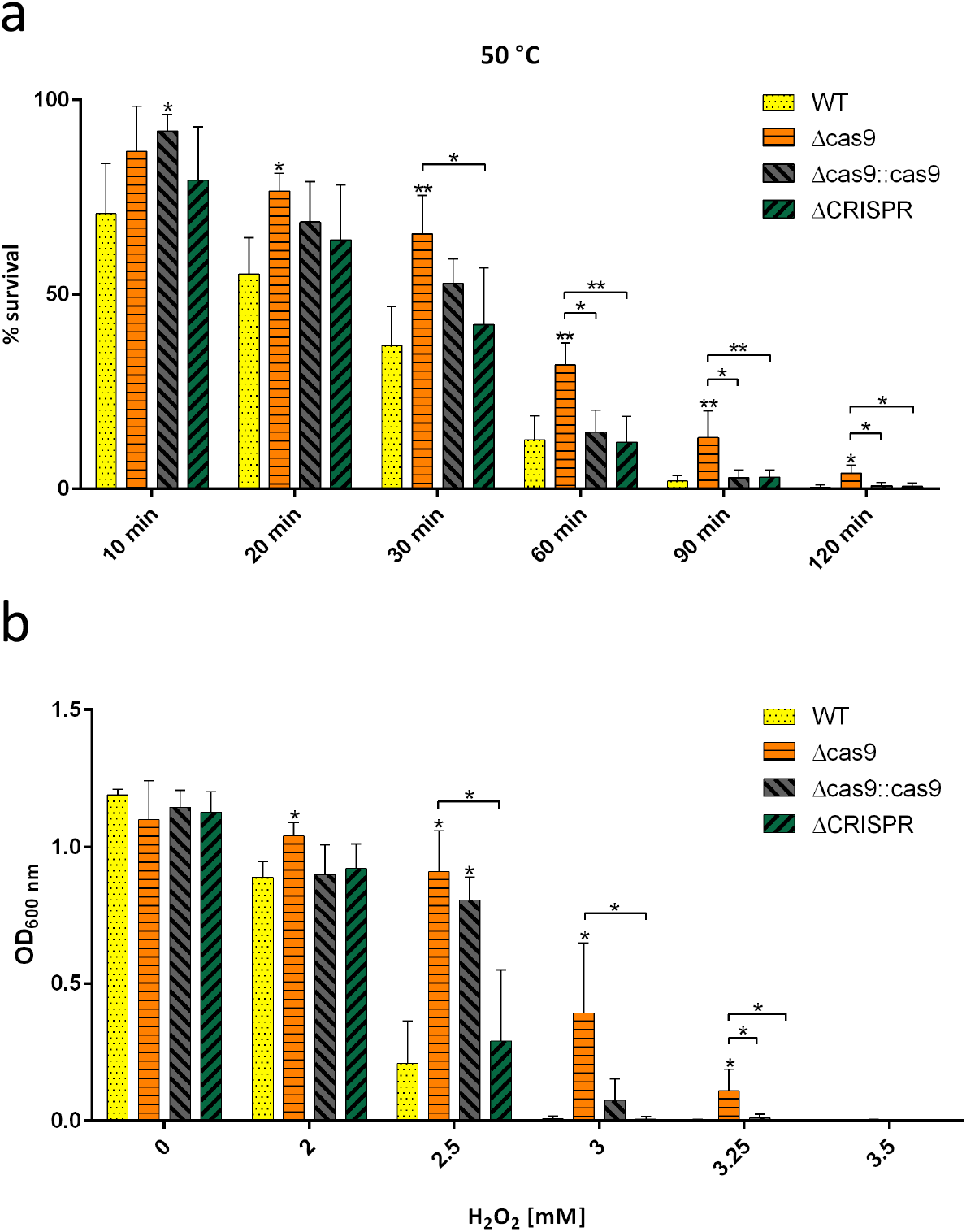
CRISPR-Cas related stress response to high temperature and oxidative stress. (a) Survival of *S. anginosus* SK52 and mutant strains after exposure to 50 °C temperature stress for 10 min – 120 min. Results represent mean CFU and standard deviations of five independent experiments conducted with the *S. anginosus* SK52 and mutant strains. (b) Effect of oxidative stress to the survival of *S. anginosus* SK52 and mutant strains. *S. anginosus* SK52, as well as Δ*cas9*, Δ*cas9::cas9* and ΔCRISPR strains were subjected to 2 mM – 3.5 mM hydrogen peroxide (H_2_O_2_) for 24 h. Results were obtained during four independent experiments and represent mean values and standard deviations. Differences were statistically significant (*p < 0.05, **p < 0.01; Mann-Whitney-U test), unless otherwise indicated it refers to wildtype strain.

Influence of Cas9 on growth behavior of *S. anginosus*

Carrying accessory genes can be accompanied by fitness costs (45). To determine whether the presence or absence of *cas9* or other mutations of the CRISPR-Cas system has an effect on growth in our experimental settings, we monitored cell densities of *S. anginosus* SK52 (WT) and the mutants Δ*cas9*, ΔCRISPR and Δ*cas9::cas9* over 24 h (Figure 4B) in liquid broth. Interestingly, the mutants Δ*cas9* and Δ*cas9::cas9* grew faster compared to WT and the ΔCRISPR mutant, while the *cas9* deficient mutant showed a more pronounced increase than the *cas9* complementation strain after 5 h, suggesting that *cas9* expression indeed is associated with the growth rate. This was further supported by comparing the colony sizes of the mentioned strains after growth of 24 h on agar plates (Figure 4A). The largest colonies on solid media were clearly formed by the *cas9* deficient strain, whereas the ΔCRISPR, Δ*cas9::cas9* and the wildtype were comparable in size.

**Figure 4:**
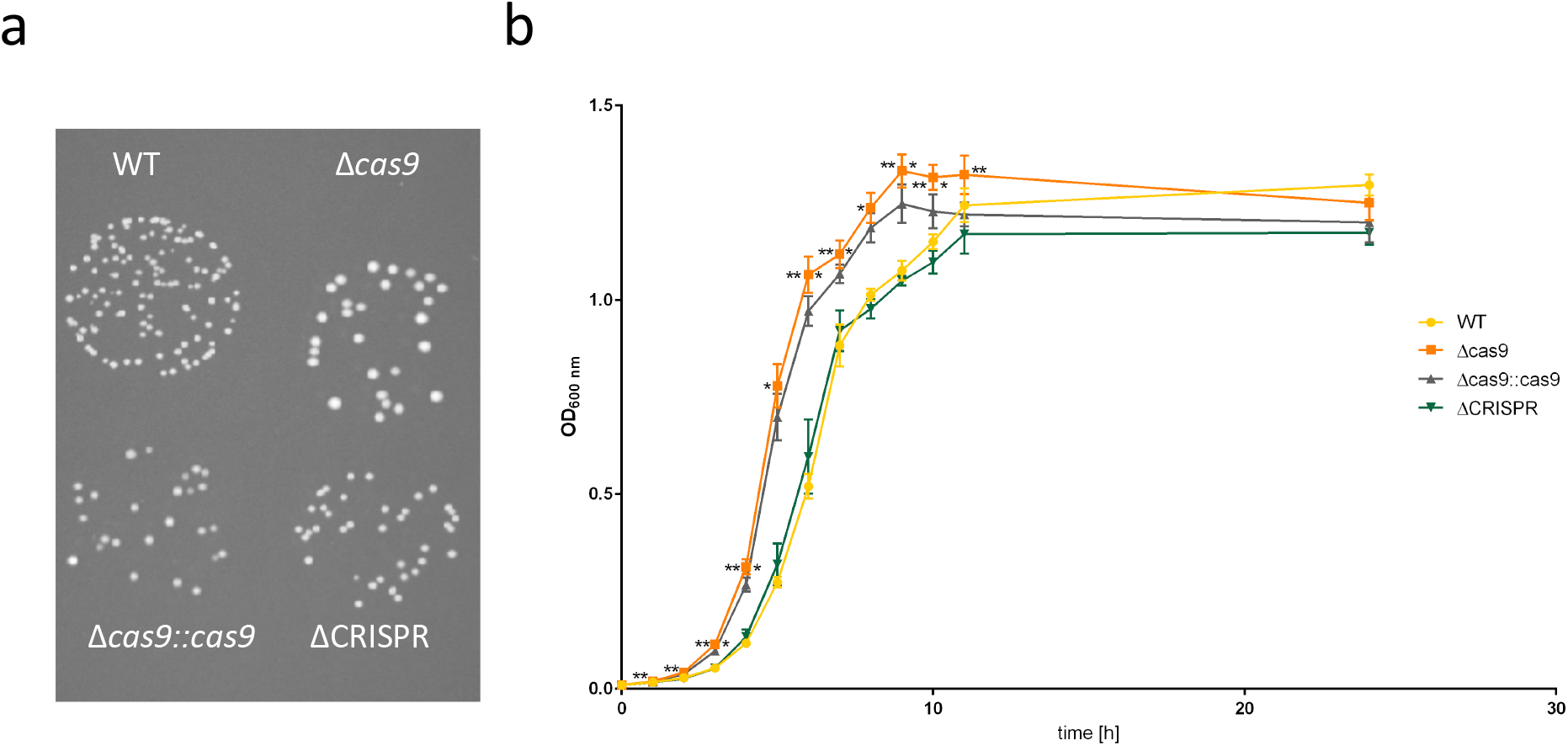
Growth behavior of *S. anginosus* influenced by Cas9. (a) Differences in colony size of *S. anginosus* SK52 (WT), *cas9* deletion mutant (Δ*cas9*), *cas9* complementation mutant (Δ*cas9::cas9*) and CRISPR array deletion mutant (ΔCRISPR). 10 µl droplets of indicated strains were incubated for 24 h on THY agar. (b) Growth of *S. anginosus* SK52 wildtype and mutant strains over time. Each point represents the average of five independent optical density values per sample including standard deviation. Differences were statistically significant (*p < 0.05, **p < 0.01; Mann-Whitney-U test). When indicated on left side of the Δ*cas9* growth curve it refers to wildtype strain and on right side it refers to Δ*cas9::cas9*.

### Metabolic activity is linked to the presence of Cas9

The results of monitoring growth in liquid media pointed to a potential metabolic burden associated with the presence of Cas9 in *S. anginosus*. To further substantiate this assumption metabolic activity of the WT and the mutant strains Δ*cas9*, ΔCRISPR and Δ*cas9::cas9* was determined by a resazurin assay (Figure 5). The measurable reduction of resazurin to resofurin by NADH or other reducing agents is commonly used as an indicator for metabolic activity in living bacterial cells (46). The absorbance measurements reflecting metabolic activity that were obtained with and without environmental stress through high temperature of 50 °C revealed significantly higher values in the *cas9* deletion strain in both settings. Overall, these results support the idea that Cas9 in *S. anginosus* is associated with higher metabolic costs.

**Figure 5:**
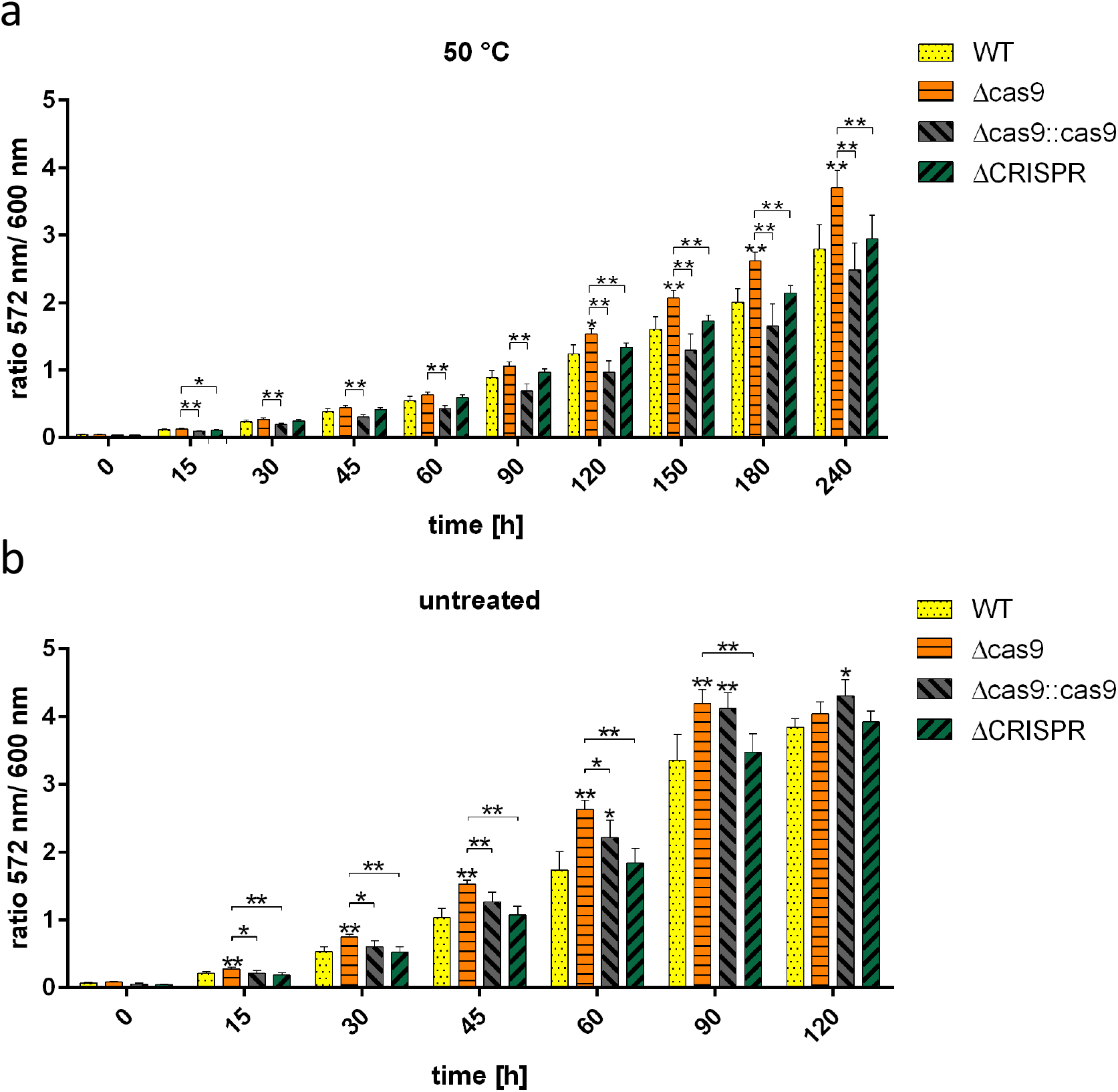
Metabolic activity related to the expression of *cas9* with and without environmental stress. (a) Effect of 50 °C temperature stress on the metabolic activity of *S. anginosus* SK52 and mutant strains within 4 h in a resazurin assay. (b) Measurement of the metabolic activity of *S. anginosus* SK52 and mutant strains within 2 h in a resazurin assay. Results represent the mean absorbance ratio 572 nm/ 600 nm and standard deviations of five independent experiments conducted with the *S. anginosus* SK52 and mutant strains. Differences were statistically significant (*p < 0.05, **p < 0.01; Mann-Whitney-U test), unless otherwise indicated it refers to wildtype strain.

## Discussion

CRISPR-Cas systems protect bacteria against harmful invading genetic elements, that are potentially threatening the bacterial cells (1). The majority of *S. anginosus* isolates harbors type II-A CRISPR-Cas systems with the prominent effector protein Cas9 (4,43). We previously reported on the diversity of *S. anginosus* CRISPR-Cas type II-A systems (16). However, the functionality of *S. anginosus* CRISPR-Cas system has not yet been investigated. This study demonstrates, that the type II-A system of *S. anginosus* is a functional adaptive immunity system through a plasmid interference assay (Figure 1B). Upon introduction of a plasmid containing an endogenous spacer along with a putative streptococcal PAM, Cas9 could efficiently cleave the target DNA in presence of a TGG sequence and therefore prevent or highly reduce plasmid transformation. This interference was abolished by the deletion of *cas9* or the entire CRISPR array as well as in presence of a random sequence instead a functional PAM. Selection of the nucleotides TGG as PAM was made based on functional PAM sequences in *S. pyogenes* (47) and confirms that PAMs of streptococcal type II-A systems work with different types of Cas9 nucleases. Efficient cleavage of the target DNA could be restored by complementing the Cas9 deletion (Figure 1B). To the best of our knowledge these data demonstrate for the first time the presence of a functional CRISPR-Cas system in the species *S. anginosus*.

CRISPR-Cas systems are also involved in processes going beyond bacterial defense, such as the regulation of bacterial physiology or virulence and the adaptation to environmental stress (21,24,48). The ability to adapt to stress is crucial for human pathogens as *S. anginosus*, as they are exposed to a variety of different microenvironments during colonization and invasion of host tissue (21). CRISPR-Cas systems are known to be activated as response to stress (23,24) and may therefore contribute to bacterial survival under harsh environmental conditions. To investigate how the stress response of *S. anginosus* is affected by CRISPR-Cas we exposed the WT and several mutants to various stress conditions. Unexpectedly however, survival under stress generated through UV-light, hydrogen peroxide and heat was significantly lower in strains carrying the *cas9* gene (Figure 2 and Figure 3). Similar to our observations, a recently published study could show that deletion of the archaeal Cas3 in *Haloferax volacanii* led to a higher resistance towards UV exposure (49).

In general, the presence of CRISPR-Cas often seems to be associated with fitness benefits as these systems can be found in different species from various environments (18) and provide protection against invasion of foreign genetic material. Nevertheless, half of all bacteria do not harbor CRISPR-Cas systems and even in entire species as *S. pneumoniae* they are absent (7,18,19). In terms of protecting themselves from invading phages, bacteria can rely on other strategies besides the adaptive immunity conferred by CRISPR such as receptor or cell-wall modifications and the restriction-modification system (50). However, as the restriction-modification system is associated with fitness costs, any system that provides resistance or defense against outer threats may be associated with costs for the host bacterium (51,52). A recent review details potential fitness costs of CRISPR-Cas and the responsible mechanism (53). Whether the presence or absence of a CRISPR-Cas system is associated with increased fitness costs may however be species and strain specific. The results of a recent study from Westra *et al.* (54) suggested that deletion of a single *cas* gene in *P. aeruginosa* had no significant effect on its ability to compete with the wildtype strain. However, the deletion of the entire type I-F CRISPR locus led to decreased fitness, indicating that carrying CRISPR is not cost-intensive for *Pseudomonas aeruginosa* and that it has additional beneficial functions apart from phage defense. For *Streptococcus thermophilus* different observations were made (52). It has been shown that maintaining CRISPR immunity systems can affect growth. In the mentioned study Cas protein expression was confirmed as particularly costly, as Cas deficient mutants were able to outgrow the wildtype strain. Leading to the conclusion that the maintenance of the adaptive immune system CRISPR-Cas is responsible for the fitness costs.

For *S. anginosus* our findings indicate that the expression of a functional CRISPR-Cas system may also be associated with fitness costs, an interpretation that is supported by the results of our growth experiments (Figure 4). Deletion of *cas9* led to a significantly enhanced growth behavior in liquid culture and the detected CFU were enlarged compared to wildtype *S. anginosus*. Complementation of *cas9* also resulted in a significantly increased growth rate compared to the wildtype but was slower than the *cas9* deficient mutant. These observations were consistent with the *cas9* transcript level (Figure 1A), where the expression of *cas9* was reduced in a *cas9* complementation strain compared to the wildtype strain and the CRISPR array deficient strain. The constant levels of *cas9* expressions that we observed in our reporter gene experiments may be responsible for the reduced growth and impaired stress survival through metabolic costs associated with Cas9 production. This interpretation is supported by resazurin assay data, which clearly show a reduced metabolic activity of strains carrying the *cas9* gene (Figure 5). So far, rather little attention has been paid to the metabolic costs related to maintenance of CRISPR-Cas. While the loss of CRISPR-Cas carries clear disadvantages regarding immunity functions, it could very well be advantageous considering bacterial metabolic functions. This trade-off may play a bigger role in bacterial species that are difficult to cultivate, such as certain streptococci. For the interpretation of our results the question arose, whether the increased survival of *cas9* deletion mutants is linked to the nuclease function of Cas9 and the loss of genetic remodeling capabilities. The Cas9 nuclease function depends of course on the presence of the CRISPR array, as confirmed with our plasmid interference assay. And since in our experiments the CRISPR array deletion mutant displayed sensitivities towards environmental stress comparable to wildtype *S. anginosus* (Figure 2 and Figure 3), we must assume that the specific nuclease activity, that is also absent in the CRISPR array deletion mutant is not responsible for an altered stress adaptation.

In theory it may be beneficial to tightly regulate the expression of CRISPR-Cas systems, due to the permanent metabolic costs of constitutive expression (54–56). Although *cas9* expression in *S. anginosus* (Figure 1A) was clearly demonstrated, no growth phase or condition could be defined in which *cas9* expression was significantly enhanced (Figure S2 & Figure S3), suggesting a constitutively expressed CRISPR-Cas type II-A system in *S. anginosus*. These findings are supported by studies confirming constitutive CRISPR-Cas expression to rapidly target invading nucleic acids (57,58) and could also be shown in *Haloferax volcanii* an archaeal species (59). While this is associated with metabolic costs, it offers an immediate benefit in terms of response time (55). For lactobacilli it is shown that their type II-A CRISPR-Cas systems must be constitutively active and ready for immediate defense as they live in a competitive environment and are therefore constantly threatened by invading DNA (58). As lactobacilli, *S. anginosus* belongs to the order of Lactobacillales, and is present in similar areas of the human body such as the oral cavity, the gastrointestinal and the urogenital tract (25–28,60).

Another point to consider is that CRISPR-Cas systems have been shown to limit the uptake of not only harmful but rather beneficial traits encoded on foreign nucleic acid that could promote fitness by increasing resistance or pathogenicity of the host bacterium (12,19,61). For pathogenic enterococci a correlation could be shown between the absence of CRISPR-Cas loci and the accumulation of antibiotic resistance genes (62). In a previous study we elucidated an inverse relationship between the presence of hemolysin genes and the carriage of CRISPR-Cas systems in *S. anginosus* (40). The presence of CRISPR in a particular *S. anginosus* strain could therefore be the result of a trade-off between the need of an adaptive defense against threatening foreign nucleic acid and fitness costs due to the metabolic load of having a constitutively active CRISPR-Cas system.

In summary we could demonstrate the presence of a functional CRISPR-Cas system in *S. anginosus,* leading to measurable metabolic costs for its maintenance. Carrying Cas9 was associated with an impaired stress response and may thus explain, why many strains of this species lack CRISPR-Cas.

## Material and Methods

### Bacterial Strains and Growth Conditions

For cultivation of *Streptococcus anginosus* strains a liquid culture in THY medium [Todd-Hewitt Broth (Oxoid) supplemented with 0.5 % yeast extract (BD)] was prepared, whereas the cultivation on solid media was conducted on sheep blood agar plates (Oxoid). Liquid cultures as well as plates were incubated at 37 °C and 5 % CO_2_. Streptococcal mutants were incubated with 120 µg ml^-1^ spectinomycin (pAT28 or pGA14) or 10 µg ml^-1^ erythromycin (pAT18 or pG^+^host5) if necessary. Lysogeny Broth (LB-Miller) was used to cultivate *Escherichia coli* DH5α and EC101. Liquid *E. coli* cultures were incubated aerobically at 37 °C on a shaker (180 rpm), while plates were kept at 37 °C with 5 % CO_2_. Four plasmids, pAT18, pAT28, pGA14 and pG^+^host5 were used in this study. 100 μg ml^−1^ spectinomycin (pAT28 and pGA14) or 400 μg mg^−1^ erythromycin (pAT18 and pG^+^host5) were supplemented for cultivation of *E. coli* strains carrying these plasmids. All strains and plasmids used in this study are listed in Table 1.

**Table 1:**
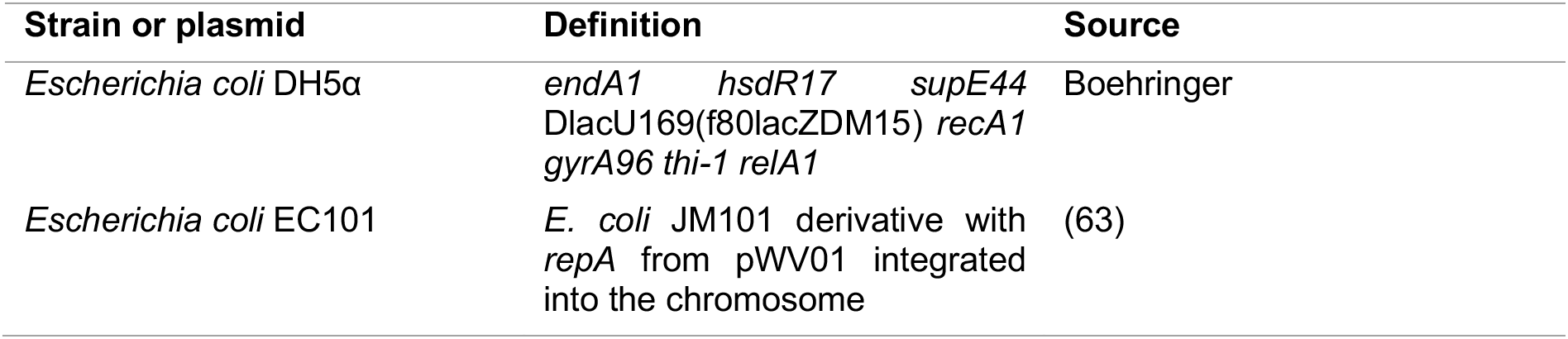
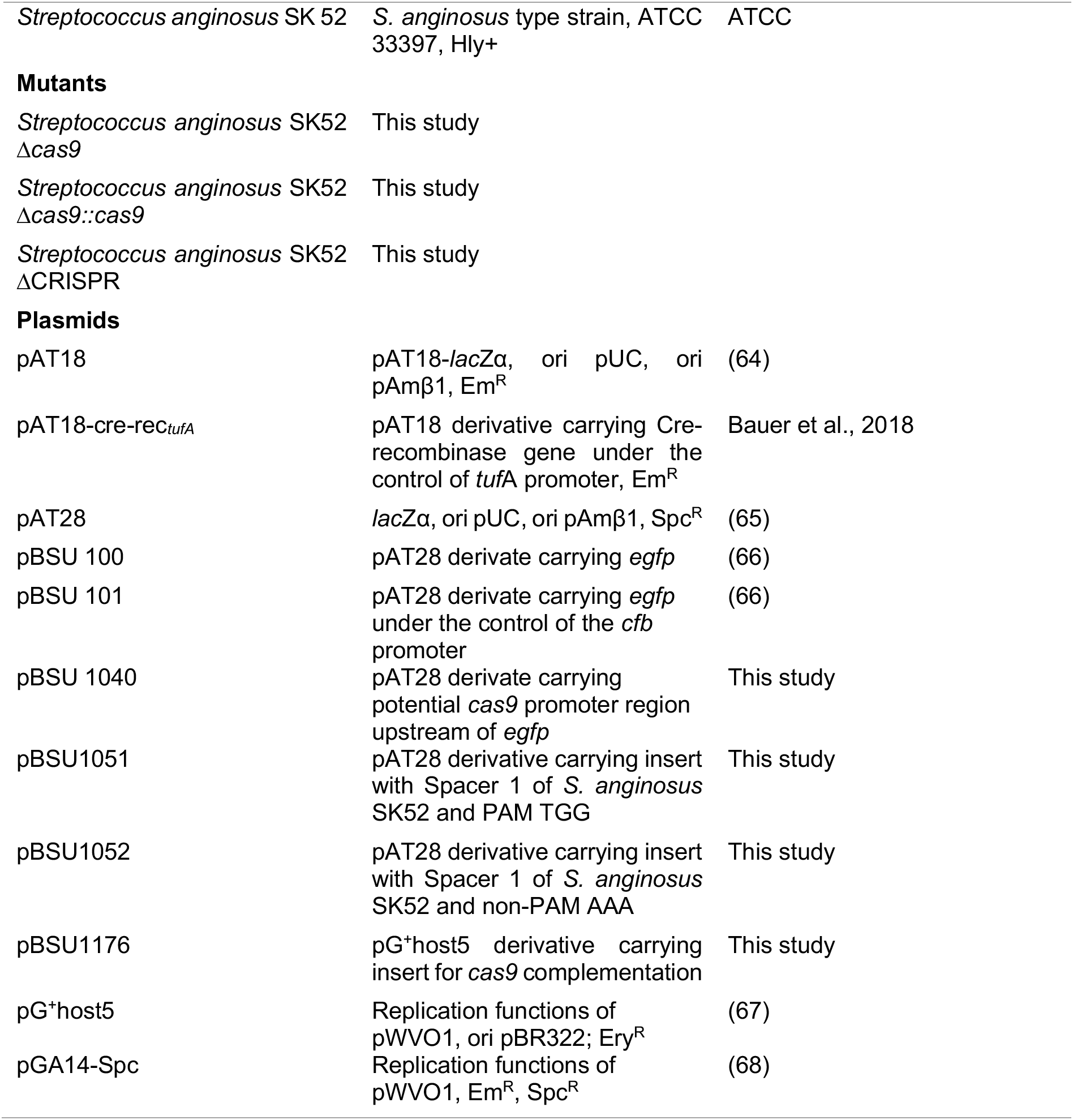
Bacterial strains, generated mutants and plasmids used for this study.

### General DNA Techniques

The isolation of DNA (GeneElute^TM^ Bacterial Genomic DNA, Sigmal-Aldrich) or plasmids (QIAprep^®^ Spin Miniprep Kit, Qiagen) was performed applying commercial kits, following the manufactureŕs protocols. DNA and plasmid concentrations were measured using Quant-iT dsDNA Broad-Range (BR) Assay Kit (Invitrogen). Polymerase chain reactions (PCR) were performed using standard protocols for the Expand Long Template PCR system: DNA pol. Mix (Roche) with buffer 3 or the *Taq* polymerase (Roche). The initial denaturation for *Taq* polymerase at 95 °C for 5 min was followed by 30 amplification cycles of 1 min at 95 °C, 30 sec at 50 °C, 1-4 min at 72 °C and a final elongation of 7 min at 72 °C, where the elongation time was adjusted according to the product length (1 min per 1000 bp). For templates longer than 3000 bp the Expand Long Template PCR system: DNA pol. Mix was applied. Default settings were an initial denaturation for 5 min at 92 °C followed by 11 cycles of 92 °C for 10 sec, 50 °C for 30 sec and 68 °C for 3 min. Followed by 25 cycles of 92 °C for 15 sec, 50 °C for 30 sec and 68 °C for 3 min + 25 sec per cycle and a final elongation of 68 °C for 5 min. Annealing temperature and the elongation time varied depending on predicted primer annealing temperature and product length. To clean up PCR products, NucleoSpin^®^ Gel and PCR Clean-up (Machery-Nagel) was used. Primers used in this study are listed in Table S1. Commercial nucleotide sequencing was carried out by Microsynth Seqlab laboratories (Göttingen, Germany).

RNA extraction for reverse transcriptase PCR (RT-PCR) was carried out using Qiagen RNeasy^®^ Mini Kit according to the manufacturers protocol with following modifications. Bacteria grown to an optical density at 600 nm (OD_600nm_) of 0.6 were pelleted and resuspended in THY broth. Bacterial suspensions were mixed with Qiagen RNAprotect^®^ Bacterial Reagent followed by 5 min incubation at room temperature. Pellets were resuspended in RLT buffer supplemented with β-Mercaptoethanol before bacterial cells were lysed with a Ribolyser at 6000 rpm for 20 sec. After centrifugation, the supernatant was transferred to Qiagen QIAshredder columns and the manufacturers protocol was continued. The isolated RNA was subjected to a DNase digestion followed by RNA concentration measurement using Qubit^®^ RNA HS system. Further, the RT-PCR was conducted with 20 ng RNA using Primers 13/41 and the Qiagen^®^ OneStep RT-PCR Kit following the instructions of the manufacturer.

## Construction of Mutants

### Construction of *cas9* and CRISPR-array deletion strains

*S. anginosus* SK52 Δ*cas9* was created via splicing by overlap extension PCR (SOE) and competence related transformation as described in Bauer et al., 2018. In brief, *cas9* flanking regions of *S. anginosus* SK52 were amplified with primer pairs 1/2 for Fragment 1 (F1) and 3/4 for Fragment 2 (F2). The primers introduced an overlap to either a *lox66* or a *lox71* site. The spectinomycin resistance gene was amplified in two fragments from pGA14-Spc using primers 5/8 and 6/7, that introduced a *lox66* or *lox71* site, respectively, adjacent to the spectinomycin gene. All fragments were fused together by SOE-PCR. The size of the fusion construct was controlled by an agarose gel (1%). Subsequently, the linear DNA construct was transformed into *S. anginosus* SK52 by inducing natural competence with competence-stimulating peptide 1 (CSP-1). SK52 was incubated with 100 ng CSP-1 and after 40 min the DNA construct was added. After two hours of incubation the bacteria were plated on THY-spectinomycin agar. Colonies were picked after 24-48 h and screened for successful insertion by colony-PCR. Therefore, streptococci were resuspended in 50 µl Aqua bidest and then incubated at 95 °C for 30 min. Subsequently, supernatant was used in a standard *Taq*-PCR (see section ‘General DNA Technique’) with primers 9/10. To eliminate the spectinomycin resistance gene, positive clones were transformed with a Cre-recombinase encoding plasmid (pAT19-cre-rec_tufA_). Transformed bacteria were plated on sheep blood agar supplemented with erythromycin. The two lox-sites are recombined by the Cre-recombinase to a singular *lox72* site. Spectinomycin-sensitive but erythromycin resistant clones were incubated without antibiotics to induce plasmid loss, resulting in a markerless deletion strain. The newly created deletion site was amplified by PCR with primers 9/10 and checked by DNA sequencing.

The same method was applied to create a markerless CRISPR array knockout strain of *S. anginosus* SK52. Primers 25/26 (F1) and 27/28 (F2) were used to generate flanking fragments and with primers 7/24 and 8/23 the lox-spectinomycin fragment was created. To verify correct deletion, primers 29/30 were used.

### Construction of *cas9* complementation strain

The temperature sensitive pG^+^host5 vector was used to create the *S. anginosus* SK52 Δ*cas9::cas9* by a chromosomal integration of *cas9* at its native locus. First, a silent point mutation was introduced into the *cas9* gene to be able to distinguish the complemented strain and the type strain. Therefore, two PCR products (F1: Primers 11/12, F2: Primers 13/14) were fused in an SOE-PCR using Primer 11/14, which additionally introduced *Xho*I and *Hind*III restriction cutting sites. The resulting linear DNA-fragment was subsequently cloned into the pG^+^host5 plasmid [digested with *Xho*I and *Hind*III (New England Biolabs)]. The resulting complementation vector was transformed into *E. coli* EC101 by heat shock transformation. Plasmid DNA was extracted and used for transformation into the *S. anginosus* SK52 using the *S. anginosus* specific competence stimulation peptide CSP-1 as described previously (41) and the regeneration of the transformants was performed for 3 h at 39 °C. Bacteria were plated on sheep blood agar supplemented with 10 µg ml^-1^ erythromycin and incubated at 37°C for 48 h. Clones resistant towards erythromycin were tested via colony-PCR for the integration of the plasmid at the desired locus with Primers 15/16. To induce loss of the chromosomal integration of pG^+^host5, cultures were grown at 30 °C. Subsequently single clones were tested for erythromycin susceptibility and the complementation of the *cas9* gene was verified by PCR (Primers 17/18, 19/20 and 21/22) and sequencing.

## Construction of plasmids

### CRISPR-Cas functionality assessing plasmids

Based on the plasmid pAT28 two different inserts were created and cloned into *E. coli* EC101. The inserts were designed to contain a single endogenous spacer sequence located in the CRISPR array of *S. anginosus* SK52, Spacer 1 and a potential functional or an intentional non-functional PAM. Inserts were constructed by annealing of oligonucleotides 31/32 and 33/34 and subsequently cloned into plasmid backbone pAT28 [both digested with *Eco*RI and *Bam*HI (New England Biolabs)]. Single colonies resistant towards spectinomycin were tested via colony-PCR with Primers 35/36 for insert presence and verified by sequencing.

### Promoter reporter plasmids

To analyze *cas9* expression a reporter construct with EGFP was created. The putative *cas9* promoter was amplified using primers 37/38 additionally introducing restriction cutting sites for *Bam*HI and *Eco*RI. Both enzymes (New England Biolabs) were used to digest both, the PCR product as well as plasmid pBSU100. After purification, both products were ligated and transformed into *E. coli* DH5α by heat shock transformation. Extracted plasmid DNA was used for transformation into *S. anginosus* strain SK52 by inducing natural competence as previously described (41). Correct plasmid construction was verified by PCR using primers 39/40 followed by DNA sequencing.

### Measurement of growth behavior

The OD_600 nm_ of overnight cultures of *S. anginosus* SK52 was determined, adjusted to 0.01 in THY broth and incubated for 24 h at 37 °C and 5 % CO_2_. Bacterial suspensions were measured every 60 min for 11 h and again after 24 h of incubation.

The colony size was compared by plating 10 µl spots of grown *S. anginosus* cultures on THY agar plates following an overnight incubation at 37 °C and 5 % CO_2_.

### Temperature sensitivity assay

To assess the ability of *S. anginosus* SK52 to tolerate heat stress, a temperature sensitivity assay was performed. After adjusting the OD_600nm_ of overnight cultures to 0.01 in THY broth, bacterial suspensions of 400 µl were incubated at 50 °C for different periods of time (0 – 120 min). Serial dilutions of bacteria were plated on THY agar plates before incubating them overnight. CFU were counted to determine the percentage of survival of the heat-treated compared to untreated bacteria.

### UV light sensitivity assay

An irradiation assay was performed, to investigate the sensitivity of *S. anginosus* SK52 towards UV light. The OD_600 nm_ of overnight cultures was determined and adjusted to 0.02 in THY broth. Bacterial suspensions were diluted according a previously determined dilution factor to ensure all cultures forming the same number of colony forming units (CFU) when untreated. Bacteria were plated in serial dilutions on THY agar plates and subsequently irradiated with UV light for a certain period of time (43 – 96 sec). Following overnight incubation, the lethal irradiation time (< 5 CFU) was determined.

### Sensitivity towards oxidative stress

The sensitivity of *S. anginosus* SK52 towards oxidative stress was investigated using hydrogen peroxide. After the OD_600nm_ of overnight cultures was adjusted to 0.01 in THY broth, different concentrations of freshly prepared hydrogen peroxide solutions (final concentration 0 – 3.5 mM) were added to the bacterial suspension (total volume 5 ml). Following overnight incubation, the OD_600nm_ was determined.

### Plasmid interference assay

To assess CRISPR-Cas functionality in *S. anginosus*, a competence based transformation efficiency assay was conducted as described elsewhere (41). In brief, plasmids containing a single endogenous spacer from *S. anginosus* SK52 and a potential functional or intentional non-functional PAM sequence pAT28_SP1_TGG, pAT28_SP1_AAA and the empty vector pAT28 were transformed into *S. anginosus* SK52 by inducing natural competence with CSP-1. Streptococcal strains were incubated with 100 ng CSP-1 and after 40 min the plasmids were added. After two hours of incubation bacteria were plated in serial dilutions on THY agar plates with and without spectinomycin to determine the transformation efficiency (transformation efficiency [%] = [CFU transformants/total CFU] x 100).

### FACS analysis

To assess the influences of different conditions on *cas9* expression in *S. anginosus* SK52, bacteria were exposed to various conditions before flow cytometry analysis. Therefore, *cas9* promoter of *S. anginosus* SK52 was cloned in front of *egfp* to construct plasmid pBSU 1040, followed by transformation into *S. anginosus* SK52. The antimicrobial substances ampicillin (final concentrations: 0.015625 µg ml^-1^ – 0.0625 µg ml^-1^) and ciprofloxacin (final concentrations: 0.15625 µg ml^-1^ – 0.625 µg ml^-1^), hydrogen peroxide (final concentration: 0.33 mM and 0.16 mM) and glucose (final concentrations: 25 mM, 12.5 mM and 5 mM) as potential stressors were supplemented to the medium before inoculation with recombinant *S. anginosus* SK52 strains. Alteration of gene expression through plasmid and chromosomal DNA was assessed after the OD_600 nm_ of an overnight culture was adjusted to 0.05 in THY broth supplemented with spectinomycin. Following an incubation of 4h, bacterial suspensions were supplemented with either 1 µg µl^-1^ isolated DNA, 300 ng µl^-1^ CSP or both. To test the influence of different growth stages on *cas9* expression, the OD_600 nm_ of overnight cultures was adjusted to 0.2 in THY broth supplemented with spectinomycin and bacterial suspensions were again incubated for 3 h, 6 h or 9 h.

After overnight incubation with stressors, after DNA treatment or following the 3 h – 9 h incubation, bacteria were harvested by centrifugation (4 °C, 3000 x g, 10 min), washed and resuspended in sterile Dulbeccós phosphate buffered saline (DPBS, pH 7.4). The mean fluorescence of 10.000 events was determined by flow cytometer. The FACSCalibur (Becton Dickinson Immunocytometry Systems) was utilized with following instrument settings: FSC: E00, SSC: 400, FL-1: 700. As controls plasmids carrying *egfp* without promoter and *egfp* under the control of the *cfb* promoter were used.

### Resazurin assay

The metabolic activity of untreated and heat-treated *S. anginosus* SK52 was determined using alamarBlue^TM^ HS Cell Viability Reagent containing resazurin (Invitrogen). After the OD_600nm_ of overnight cultures was adjusted to 0.2 in THY broth, half of the bacterial suspension underwent heat-treatment for 1 h at 50 °C. According to manufacturer’s instructions 1/10th volume of cell viability reagent was mixed with heat-treated and untreated bacterial suspensions or the medium control. During the incubation time of 4 h at 37 °C the absorbance of reagent was monitored at 572 nm and 600 nm in a plate reader at certain timepoints.

### Bioinformatic and Statistical Analysis

The GenBank database (https://www.ncbi.nlm.nih.gov/) was used as source for nucleotide sequences. With the Basic Local Alignment Search Tool (42) homology searches were conducted (https://blast.ncbi.nlm.nih.gov/Blast.cgi). The genetic analysis was carried out using QIAGEN CLC Main Work Bench 7 (https://digitalinsights.qiagen.com/). The statistical analysis and preparation of graphs was performed by GraphPad Prism V6 (GraphPad Software Inc., La Jolla, CA, United States).

## Acknowledgments

The work of RB, DH and BS was supported through the Bausteine Program of the Medical Faculty, University of Ulm and the International Graduate School in Molecular Medicine Ulm. Special thanks goes to Anita Marchfelder, Ulm University, and the SPP2141 for fruitful and invaluable discussions.

## Supplement Figures and Tables

**Figure S1:**
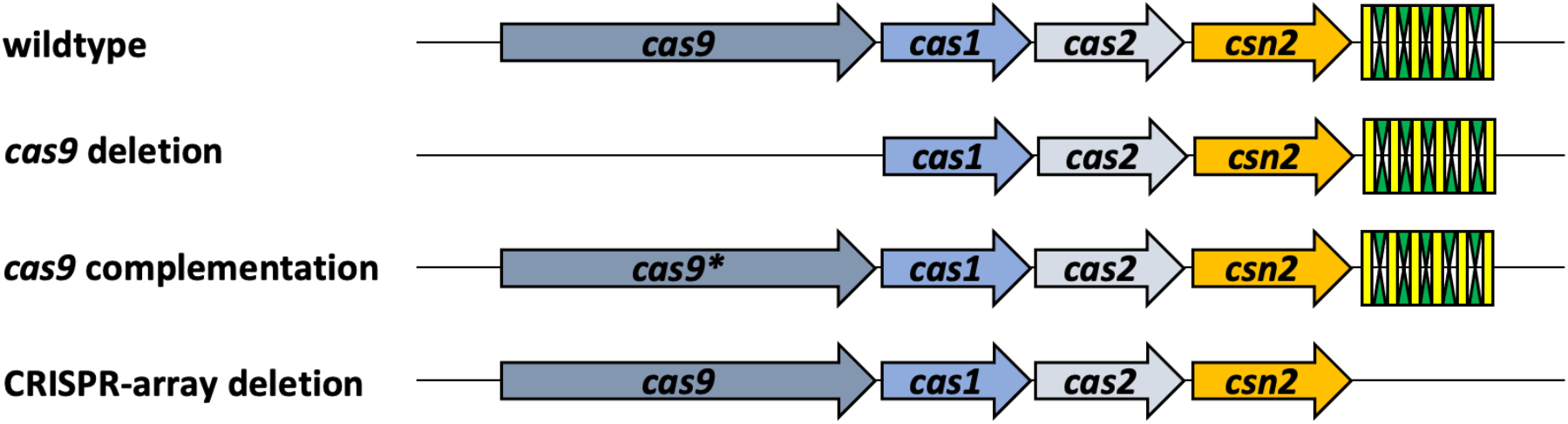
Schematic illustration of the wildtype and different mutant *S. anginosus* SK52 CRISPR-Cas type II-A loci. Arrows represent *cas* genes (dark grey: *cas9*, blue: *cas1*, light grey: *cas2* and orange *csn2*) whereas the CRISPR-array with its spacer and repeat units is shown in yellow and green (number of shown spacer and repeat units does not correspond to actual number).

**Figure S2:**
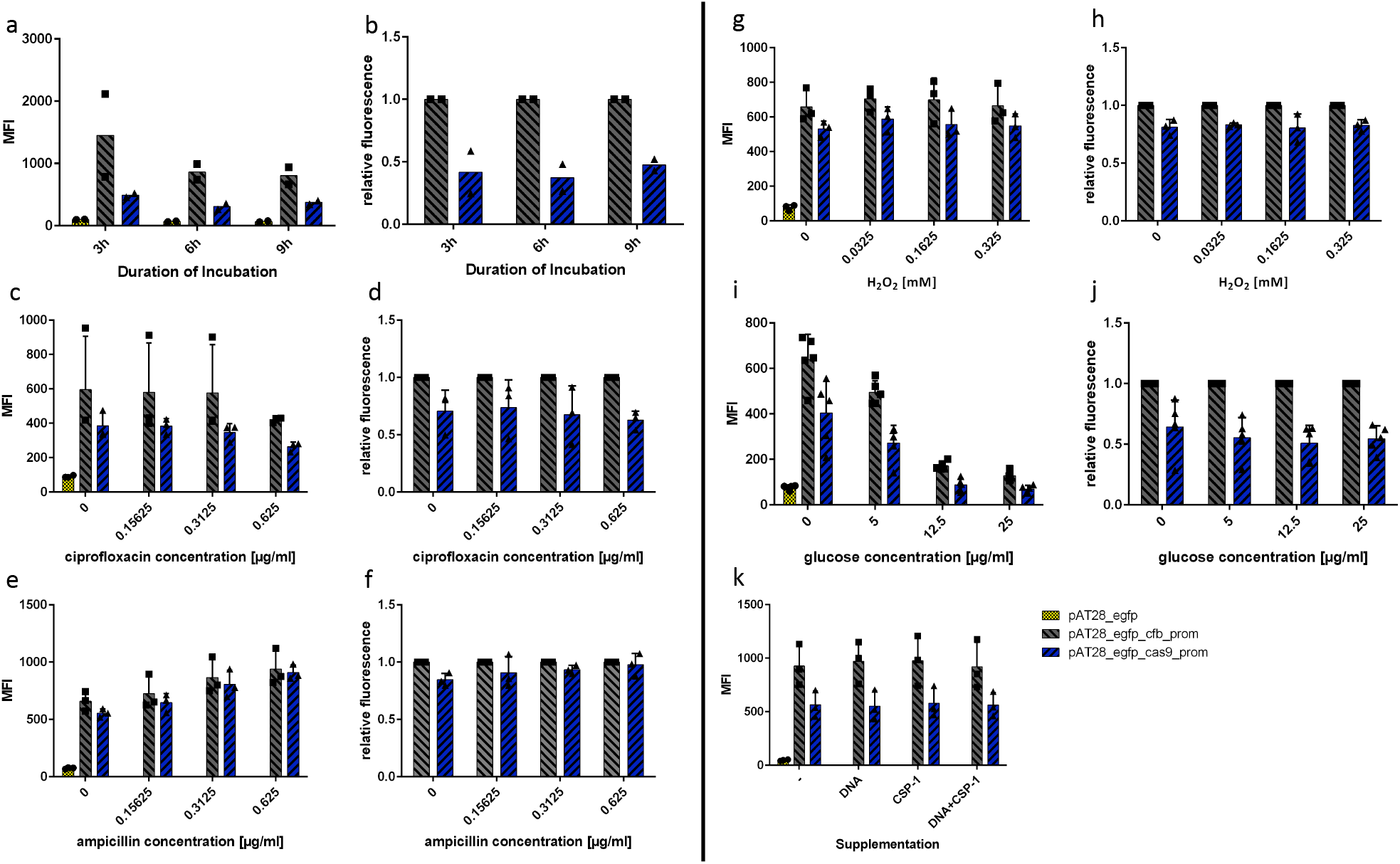
Expression of *cas9* in *S. anginosus* by measuring EGFP activity in different growth phases and under potential CRISPR-Cas inducing conditions. FACS analysis was performed for *S. anginosus* SK52 carrying reporter plasmid with *cas9* promotor (blue), the positive control carrying reporter plasmid with *cfb* promotor (grey) and the negative control carrying promotorless EGFP plasmid (yellow). Mean fluorescence intensity (MFI) corresponding to the mean value of either two, three or five probes; distribution of values is depicted. Normalization of MFI values of *S. anginosus* SK52_*cas9*_prom to the positive control *S. anginosus* SK52_*cfb*_prom. (a) Fluorescence intensity level after three, six and nine hours of initial inoculation (n=2). Normalization of MFI values in (b). (c,e) Expression of *cas9* upon supplementation with ciprofloxacin or ampicillin (n=3). Fluorescence ntensity levels measured at different concentrations (0.625 µg/ml – 0.15625 µg/ml). Normalization of MFI values in (d,f). (g) Fluorescence intensity levels recorded at different hydrogen peroxide concentrations between 0.325 mM – 0.0325 mM (n=3). Normalization of MFI values in (h). (i) MFI at different glucose concentrations between 25 mM – 5 mM (n=5). Normalization of MFI values in (j). (k) Fluorescence intensity levels measured without supplementation, for DNA supplementation of *S. anginosus* chromosomal DNA, CSP-1 supplementation (for induction of cellular DNA uptake) or supplementation of both (n=3).

**Figure S3:**
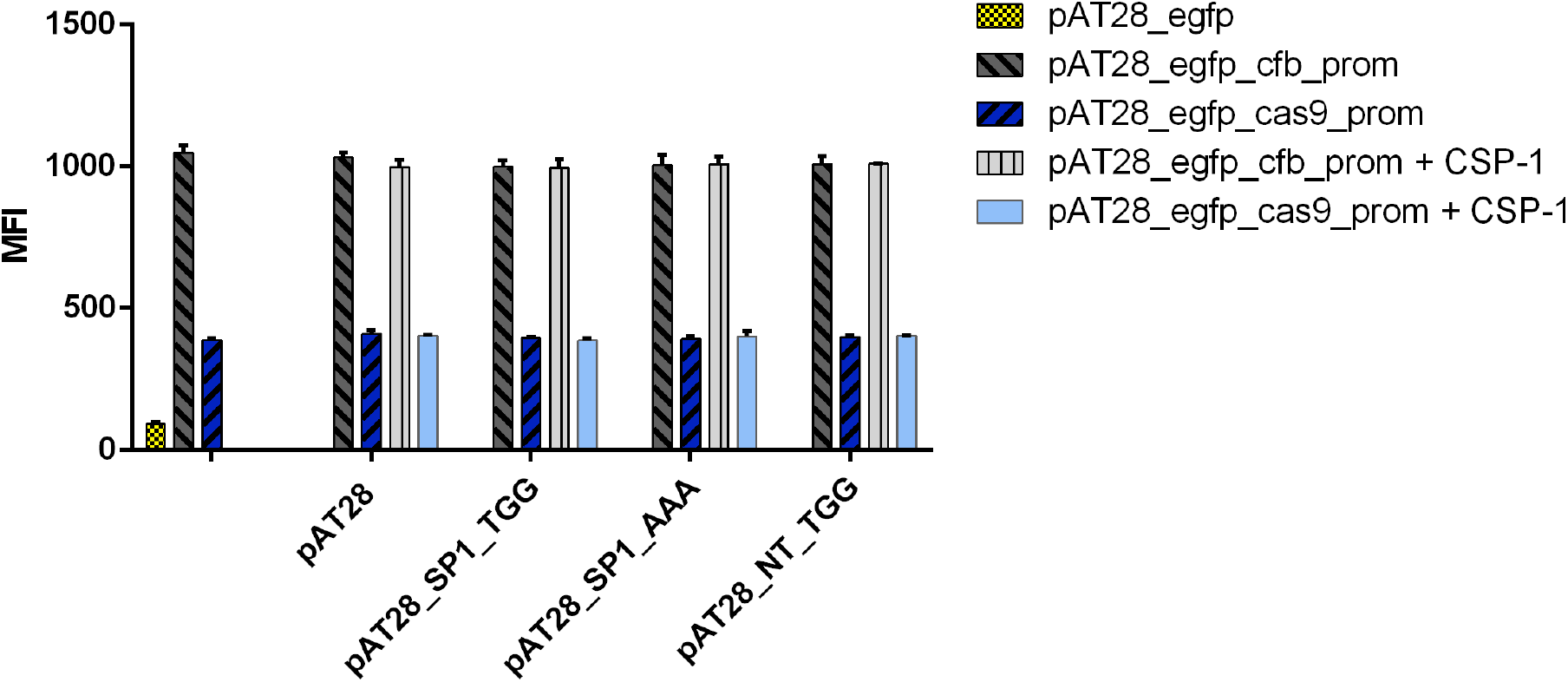
Expression of *cas9* in *S. anginosus* by measuring EGFP activity upon supplementation of plasmid DNA that induce CRISPR-Cas type II-A mediated immunity. Fluorescence intensity levels measured for supplementation of different vector constructs in presence or absence of *S. anginosus* specific CSP-1 (induction of cellular DNA uptake). Vectors were designed to contain an endogenous spacer of *S. anginosus* SK52 CRISPR-Cas sytem (SP1) or a non-target sequence (NT) followed by a functional or non-functional PAM sequence. FACS analysis was performed for *S. anginosus* SK52 carrying reporter plasmid with *cas9* promotor in presence or absence of CSP-1 (light blue and dark blue), the positive control carrying reporter plasmid with *cfb* promotor in presence or absence of CSP-1 (light grey and dark grey) and the negative control carrying promotorless EGFP plasmid (yellow). MFI corresponding to the mean value of three probes (n=3).

**Table S1:**
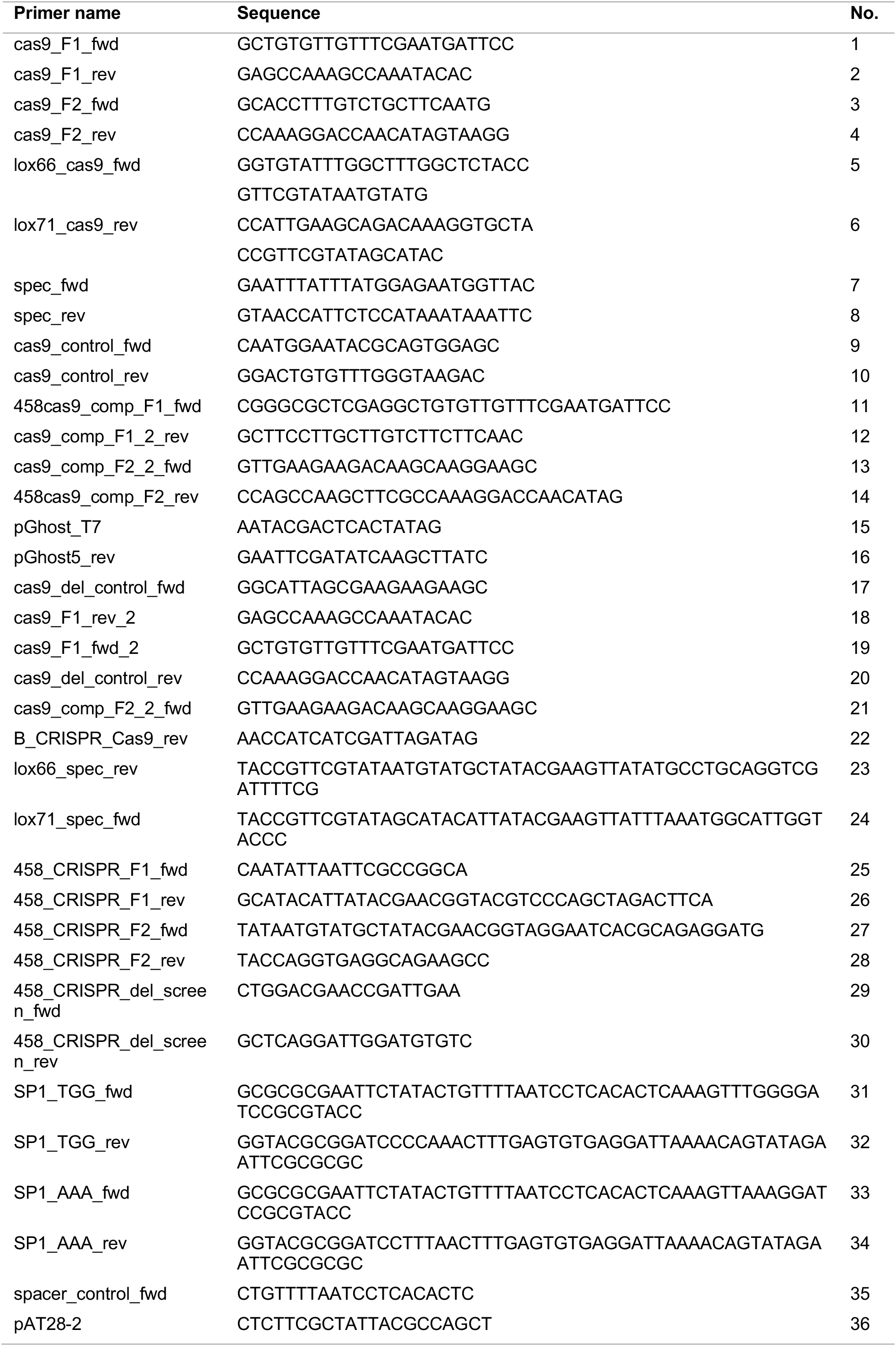
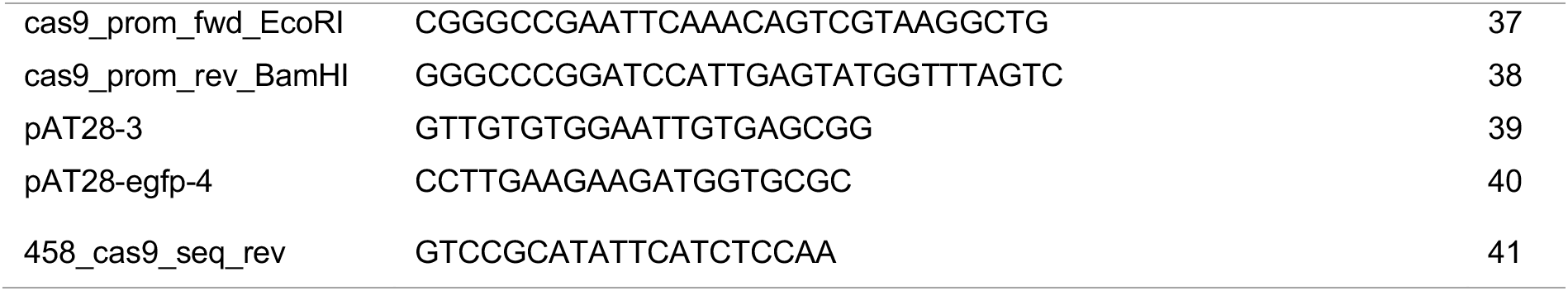
Primers used for this study.

